# Generative Deep Learning and Molecular Dynamics Reveal Design Principles for Amyloid-Like Antimicrobial Peptides

**DOI:** 10.64898/2026.03.21.713424

**Authors:** Anup K. Prasad, Vijay Awatade, Manoj K. Patel, Fabian Plisson, Lisandra L. Martin, Ajay S. Panwar

**Author notes:** Department of Biochemistry and Molecular Biology, University of Georgia, Athens, U.S.A. Department of Chemical Engineering, Indian Institute of Technology, Madras, India (Manoj K. Patel).

## Abstract

Antimicrobial peptides (AMPs) are emerging as promising alternatives to conventional antibiotics, and growing evidence indicates a fundamental link between antimicrobial activity and amyloid-like self-assembly. Many AMPs are known to form amyloid-like fibrils, while several amyloidogenic peptides exhibit intrinsic antimicrobial properties, suggesting shared underlying physicochemical determinants such as amphipathicity, β-sheet propensity, and charge distribution. However, the rational design of peptides that simultaneously encode these dual functionalities remains a significant challenge. Here, we present amyAMP, a generative deep-learning framework based on a Wasserstein generative adversarial network with gradient penalty (WGAN-GP), designed to learn and generate peptides with integrated antimicrobial and amyloidogenic properties. Trained on curated datasets of antimicrobial and amyloid-forming peptides, amyAMP captures the latent sequence-property relationships governing dual functionality. Statistical and latent-space analyses demonstrate that the generated peptides closely overlap with biologically relevant peptide space while remaining distinct from random sequences, indicating successful learning of key biochemical features. To validate functional behavior, we performed extensive coarse-grained molecular dynamics simulations to probe membrane interaction, peptide self-assembly, and membrane disruption. The simulations reveal rapid membrane adsorption, stable amphipathic insertion, and strong peptide-peptide aggregation. Notably, cooperative clustering of peptides on membrane surfaces induces membrane thinning and curvature perturbations, highlighting a mechanistic coupling between aggregation and antimicrobial activity. Collectively, these results establish that amyAMP effectively captures the shared physicochemical principles underlying antimicrobial action and amyloid-like self-assembly. This work provides a generalizable framework for the AI-guided design of multifunctional peptides to advance the development of next-generation therapeutics targeting antimicrobial resistance.

## Introduction

The global rise of antimicrobial resistance (AMR) has become a major threat to human health, with around five million deaths linked with bacterial AMR only, and projected to cause more annual deaths than cancer by 2050 if left unaddressed.^1,2^ The effectiveness of conventional antibiotics has been increasingly compromised due to the rapid evolution of resistance mechanisms in pathogenic bacteria.^3,4^ This growing crisis and social burden have created an urgent need to develop novel antimicrobial strategies that circumvent traditional resistance pathways and provide sustainable, long-term solutions.^2^ Antimicrobial peptides (AMPs) represent one of the most promising alternatives to conventional antibiotics, emerging as versatile molecular frameworks for combating multidrug-resistant pathogens.^4–6^ As evolutionarily conserved components of innate immunity, they serve as a rapid, broad-spectrum defense mechanism across all kingdoms of life.^7,8^ Their ability to disrupt microbial membranes, modulate host immune responses, and even penetrate intracellular targets provides them with multifunctional therapeutic potential.^9–11^ Moreover, AMPs exhibit exceptional sequence diversity and conformational plasticity: transitioning between α-helical, β-sheet, or disordered states depending on the environment, which underpins both their potency and adaptability.^8,12^ These attributes have made AMPs attractive scaffolds for computational and artificial intelligence (AI)-driven peptide design strategies aimed at producing next-generation antibiotics.^13,14^

Intriguingly, recent findings reveal that antimicrobial activity and amyloidogenesis are mechanistically intertwined.^15–17^ Although amyloids were historically viewed as pathological aggregates linked to neurodegenerative diseases, numerous studies now highlight their functional roles in normal physiology, including storage, scaffolding, and host defence roles.^15,17^ Several natural AMPs, such as LL-37, uperin 3.5, and temporins display an intrinsic ability to form β-rich amyloid-like fibrils that enhance membrane disruption and biofilm inhibition.^16,18–20^ These amyloid assemblies exhibit remarkable stability and protease resistance, properties that likely evolved to sustain antimicrobial function under harsh physiological or microbial conditions.^17,18,21^ The coexistence of antimicrobial potency and amyloid-like stability, thus represents an evolutionary optimization of peptide functionality, where structural self-assembly directly contributes to biological efficacy.^15–17,22^

This emerging paradigm highlights amyloidogenic AMPs as a new frontier in peptide-based drug discovery, unifying principles of self-assembly, membrane disruption, and evolutionary adaptation. Understanding the structural and energetic interplay between amyloid formation and antimicrobial action will be essential for designing next-generation therapeutics that exploit this dual functionality.^23–25^

The exploration of peptide sequence space for functional antimicrobial and amyloidogenic properties is inherently challenging due to its vastness and the resource-intensive nature of experimental screening.^25,26^ Machine learning (ML) has emerged as a powerful tool to accelerate peptide discovery by capturing complex sequence-function relationships and enabling the design of novel candidates with predefined properties.^27–29^ Generative deep learning models, including Variational Autoencoders (VAEs), Generative Adversarial Networks (GANs), and Bidirectional Generative Adversarial Networks (BiGANs), can learn distributions of biological sequences and propose new designs that are difficult to identify through conventional approaches.^27,30–33^ Among these, BiGANs provide a unique framework by simultaneously learning the generator (data synthesis) and encoder (data-to-latent mapping), allowing both sequence generation and extraction of latent physicochemical and structural features.^32,34^ Such representations facilitate the exploration of overlaps between antimicrobial and amyloidogenic peptides, guiding the generation of hybrid functional sequences.^28,30,31^

Recent studies demonstrate that ML models can effectively predict antimicrobial activity, optimize peptide properties, and even assess amyloidogenic propensity.^29,35–39^ Integrating ML with molecular dynamics simulations further enhances the ability to evaluate peptide stability, conformational dynamics, and interactions with membranes, providing a complementary computational approach to experimental validation.^40,41^ In particular, MD simulations offer mechanistic insights into peptide-membrane binding, aggregation behavior, and membrane disruption processes, which are critical determinants of antimicrobial function and are often inaccessible from sequence-based models alone. By capturing latent features associated with sequence, structure, and physicochemical properties, ML-driven generative approaches, when combined with simulation-based validation, provide a powerful strategy for the rational design of multifunctional peptides.

In this study, we present a BiGAN-based generative model for designing amyloidogenic antimicrobial peptides. By training on curated datasets of antimicrobial and amyloid-forming sequences, the model learns the defining characteristics of both functional classes and generates novel candidates spanning this dual-functional space. Computational characterization of the generated peptides reveals favorable structural and physicochemical attributes consistent with antimicrobial and amyloidogenic behavior. Furthermore, coarse-grained MD simulations are employed to evaluate peptide-membrane interactions, self-assembly propensity, and membrane disruption mechanisms, providing mechanistic validation of the predicted functionalities. Together, this integrative framework demonstrates how coupling ML-based generative design with physics-based simulations can accelerate the discovery of multifunctional peptides. More broadly, leveraging insights from functional amyloid biology offers a promising avenue for designing next-generation peptide therapeutics that exploit the synergy between amyloid stability and antimicrobial efficacy.

## 2. Methods

### 2.1 Data Collection and Preprocessing

To train the BiGAN-peps model, we constructed a curated dataset comprising peptides that exhibit both antimicrobial and amyloidogenic properties. Peptide sequences were collected from multiple databases representing two major functional classes: antimicrobial peptides (AMPs) and amyloid-forming peptides (AMYs). Antimicrobial peptide sequences were obtained from the *Database of Antimicrobial Activity and Structure of Peptides (DBAASP)*, retaining sequences between 3 and 30 amino acids in length.^42^ Entries containing non-canonical residues or duplicate sequences were removed, yielding a refined dataset of 15,938 unique AMPs. Amyloidogenic peptides were compiled from *CARs-DB*, *WALTZ-DB*, and *CPAD 2.0* databases, providing 9,502 unique sequences.^43–46^

Given the emerging evidence that certain AMPs display amyloid-like aggregation behavior and that amyloidogenic peptides can possess antimicrobial activity, we applied an additional cross-property screening to identify peptides sharing both functional attributes.^15,24,47–49^ Specifically, AMP sequences were evaluated for amyloidogenic propensity using the *WALTZ* prediction algorithm, while AMY sequences were assessed for antimicrobial potential via the *AI4AMP* deep-learning classifier.^45,50,51^ This bidirectional screening yielded 5,786 AMPs with predicted amyloidogenic signatures and 1,101 AMYs with antimicrobial features. Both subsets were merged to produce a final dataset of 6,887 hybrid-function peptides, capturing the physicochemical continuum between antimicrobial and amyloidogenic functionalities.

To ensure uniform sequence representation, peptides shorter than 30 residues were padded with a neutral placeholder residue (“X”) that carried a zero-valued vector. Each sequence was then converted into a numerical representation using the PC6 encoding scheme, which maps each amino acid to six quantitative physicochemical properties, including hydrophobicity, side-chain volume, polarity, isoelectric point, dissociation constant, and net charge index (see Table S1).^50^This encoding approach preserves both local residue features and global compositional trends, providing a robust input representation for generative modelling. The complete dataset curation and preprocessing workflow is illustrated in Figure 1.

**Figure 1.**
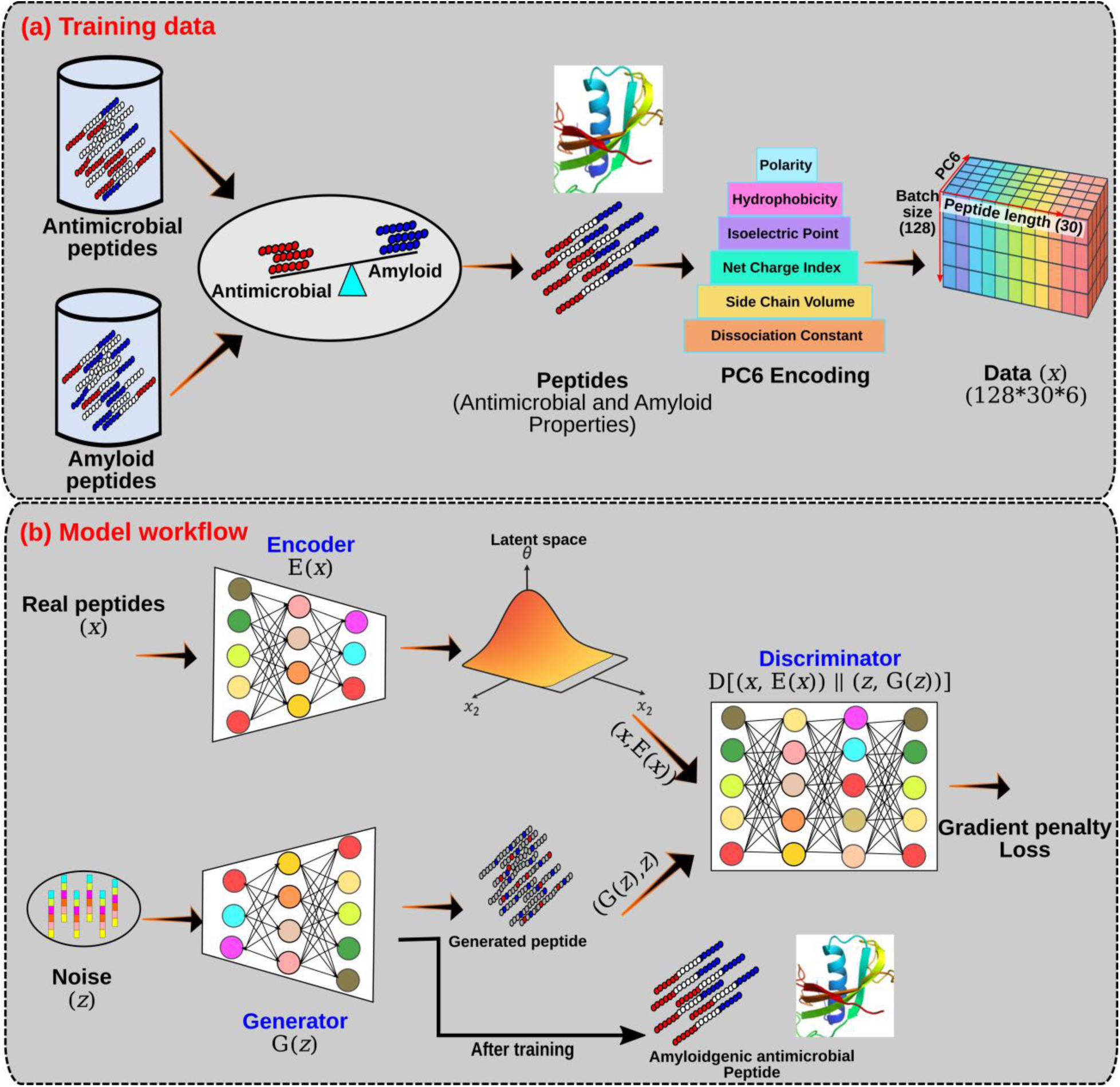
Schematic overview of the amyAMP model workflow. (a) Training data preparation: Antimicrobial and amyloid peptide datasets are collected, screened for cross-properties, and numerically encoded using the PC6 method. (b) Model framework: The amyAMP architecture integrates three core components: encoder (E), generator (G), and discriminator (D) that interact in a minimax game. The generator progressively learns to produce novel peptides that capture the statistical features of the training datasets, while the discriminator distinguishes between real and generated samples. (c) Model architecture: The internal design of the framework is depicted, emphasizing the convolutional and transposed convolutional layers used to encode and decode peptide features.

### 2.2 Model architecture

The amyAMP framework is implemented as a bidirectional generative adversarial network (BiGAN) comprising three core components: an encoder (E), a generator (G), and a critic/discriminator (C). Architectural details of these components are provided in Figure S1. Collectively, these modules learn a shared latent representation linking peptide physicochemical feature maps with their corresponding latent vectors.

#### Encoder (E)

The Encoder maps PC6-encoded peptide feature matrices into a 100-dimensional latent space. It is composed of a hierarchy of convolutional–LeakyReLU (CR) blocks, each containing a 2D convolution layer followed by a LeakyReLU activation (slope 0.2). The architecture progressively increases the depth of learned features through four stacked CR blocks with kernel sizes designed to capture residue-level and sequence-length scale features. A final convolutional layer produces a compact latent embedding, yielding:

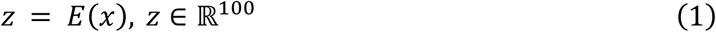

#### Generator (G)

The Generator reconstructs peptide feature maps from latent vectors using transposed convolution-BatchNorm-ReLU (TransposedCBR) blocks. Starting with a 100-dimensional latent vector, the module applies four upsampling blocks with progressively decreasing channel depth, enabling hierarchical decoding of peptide physicochemical properties. A final transposed convolution layer with Tanh activation outputs a reconstructed PC6 representation:

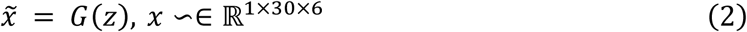

corresponding to 30-residue, 6-feature peptide matrices.

#### Critic (C)

The Critic evaluates joint pairs of peptide matrices and latent vectors, distinguishing real pairs, (𝑥, 𝐸(𝑥)), from synthetic pairs, (𝐺(𝑧), 𝑧). The critic comprises fully connected layers with LeakyReLU activations and BatchNorm, culminating in a scalar score:

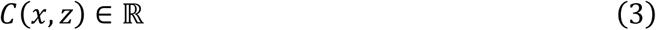

This architecture enables adversarial learning over joint data–latent distributions, as required by BiGAN theory.

#### Weight Initialization

All convolutional and linear layers were initialized with a normal distribution 𝑁(0,0.02). Batch normalization layers used 𝜇 = 1.0, 𝜎 = 0.02 for weights and zero for biases, following standard GAN initialization heuristics.

### 2.3 Training Strategy and Loss Functions

The WGAN-GP model was implemented using convolutional neural networks (CNNs), with kernel sizes adjusted according to the dimensions of the input and output data. A total of 6,887 peptide sequences were used as the training dataset, with a batch size of 128. The model was trained for 40,000 epochs, during which the dataset was repeatedly fed into the network.

To mitigate mode collapse and training instability, the Earth Mover’s (Wasserstein-1) distance was employed to measure the divergence between real and generated data distributions. In addition, a gradient penalty term was incorporated into the loss function to enforce Lipschitz continuity and stabilize training.

Training of amyAMP follows the Wasserstein GAN with Gradient Penalty (WGAN-GP) formulation, adapted for BiGANs. The objective is to minimize the Wasserstein-1 distance between the true joint distribution,

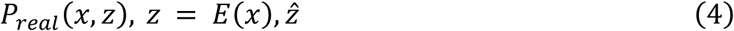

and the synthetic joint distribution,

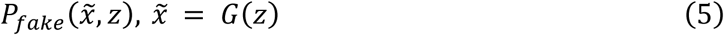

The **Critic loss** is:

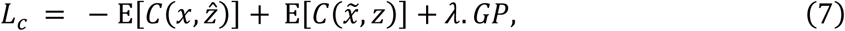

where the **gradient penalty (GP)** enforces the 1-Lipschitz constraint:

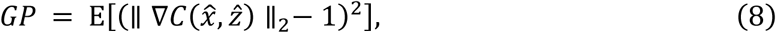

## With interpolated samples

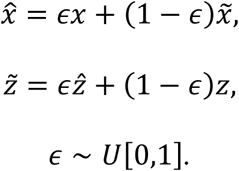

The Generator and Encoder cooperate to fool the Critic by maximizing the critic’s estimate of real pairs:

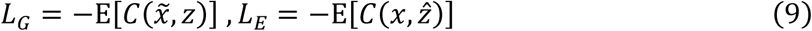

Thus, G learns to produce realistic peptide representations, while E learns to produce latent vectors consistent with the generator’s latent distribution.

### 2.4 Molecular dynamics simulations

Coarse-grained molecular dynamics (MD) simulations were performed to provide physics-based validation of the dual antimicrobial and amyloidogenic properties of peptides generated by the amyAMP model. From the library of 1000 generated sequences, 100 peptides were selected for simulation. For comparison with experimentally characterized antimicrobial peptides used during model training, reference peptides were ranked according to their reported minimum inhibitory concentration (MIC) values. From this dataset, 50 peptides spanning three activity regimes, low, intermediate, and high MIC, were selected and categorized as low-MIC, intermediate-MIC, and high-MIC peptides.

Three complementary simulation protocols were designed to probe distinct mechanistic aspects of peptide behavior:

**(i) Peptide-membrane binding simulations:** Individual peptides were simulated in the presence of a bacterial membrane model to investigate membrane association and insertion behavior. All 100 peptides were simulated for 1 µs.
**(ii) Peptide self-assembly simulations:** Systems containing eight identical peptide molecules in aqueous solution were simulated to evaluate aggregation propensity associated with amyloid-like self-assembly. All 100 peptides were simulated for 1 µs.
**(iii) Membrane disruption simulations:** Systems comprising nine identical copies of peptides interacting with bacterial membrane models were simulated to assess membrane destabilization at elevated peptide concentrations. Three representative peptides were selected for this analysis and simulated for 10 µs.

Membrane bilayers were constructed using phospholipid compositions representative of bacterial membranes. Gram-negative membranes were modelled using a 3:1 molar ratio of POPE:POPG lipids, whereas Gram-positive membranes were represented by a 4:1 POPE:POPG composition. Initial membrane configurations were generated using the CHARMM-GUI web server.^52,53^ All simulations were performed on GROMACS platform using the MARTINI coarse-grained force field with a 20 fs integration time step.^54,55^ Peptide-membrane binding and peptide self-assembly simulations were run for 1 µs, while membrane disruption simulations were extended to 10 µs to capture slower membrane remodeling events. Periodic boundary conditions were applied in all directions, and interactions were evaluated using the Verlet cutoff scheme. Electrostatic interactions were treated with the reaction-field method (dielectric constant = 15) with a cutoff distance of 1.1 nm, and van der Waals interactions were truncated at 1.1 nm using a potential-shift modifier. Temperature was maintained at 310 K using the velocity-rescale thermostat, and pressure was controlled semi-isotropically at 1 bar with the Parrinello–Rahman barostat.^56,57^ Simulation box dimensions were set to 100 × 100 × 150 nm for single-peptide membrane systems and approximately 105 × 105 × 150 nm for peptide aggregation systems.

## 3. Results and Discussion

### 3.1 Progression of learning and peptide generation

Monitoring the evolution of generated sequences during model training is critical for evaluating whether a generative model is successfully learning the statistical features of the target sequence space. Such analyses have been widely used in peptide and protein generative frameworks, including models such as PepGAN and ProteinGAN, where progressive convergence toward biologically meaningful sequence distributions serves as an indicator of effective training.^58,59^ To assess the learning progress of the amyAMP model, 1000 peptide sequences were generated at intervals of 100 training epochs and compared against the training datasets. Although each peptide in the curated training set possesses both antimicrobial and amyloidogenic characteristics, the learning statistics were analysed independently against the AMP and AMY datasets to more rigorously evaluate whether the model captures sequence patterns associated with each functional property.

Sequence similarity between generated peptides and the training datasets was quantified using percent identity. As shown in Figure 2 (a-c), the generated sequences exhibited clear evidence of progressive learning throughout training. At early epochs, generated peptides showed approximately 45% identity relative to the training datasets. As training progressed, this similarity gradually increased, reaching 52.44% and 50.99% identity relative to the AMP and AMY datasets, respectively. Importantly, when compared with a control dataset of randomly generated peptides (RandPep), the amyAMP-generated sequences maintained a stable and low percent identity throughout the training process. This observation indicates that the increase in similarity to the AMP and AMY datasets reflects targeted learning of biologically relevant sequence patterns rather than nonspecific convergence toward random peptide space. Training was therefore continued until convergence of these metrics was observed, reaching a steady state at approximately 40,000 epochs, beyond which no substantial improvement in sequence similarity was detected.

**Figure 2.**
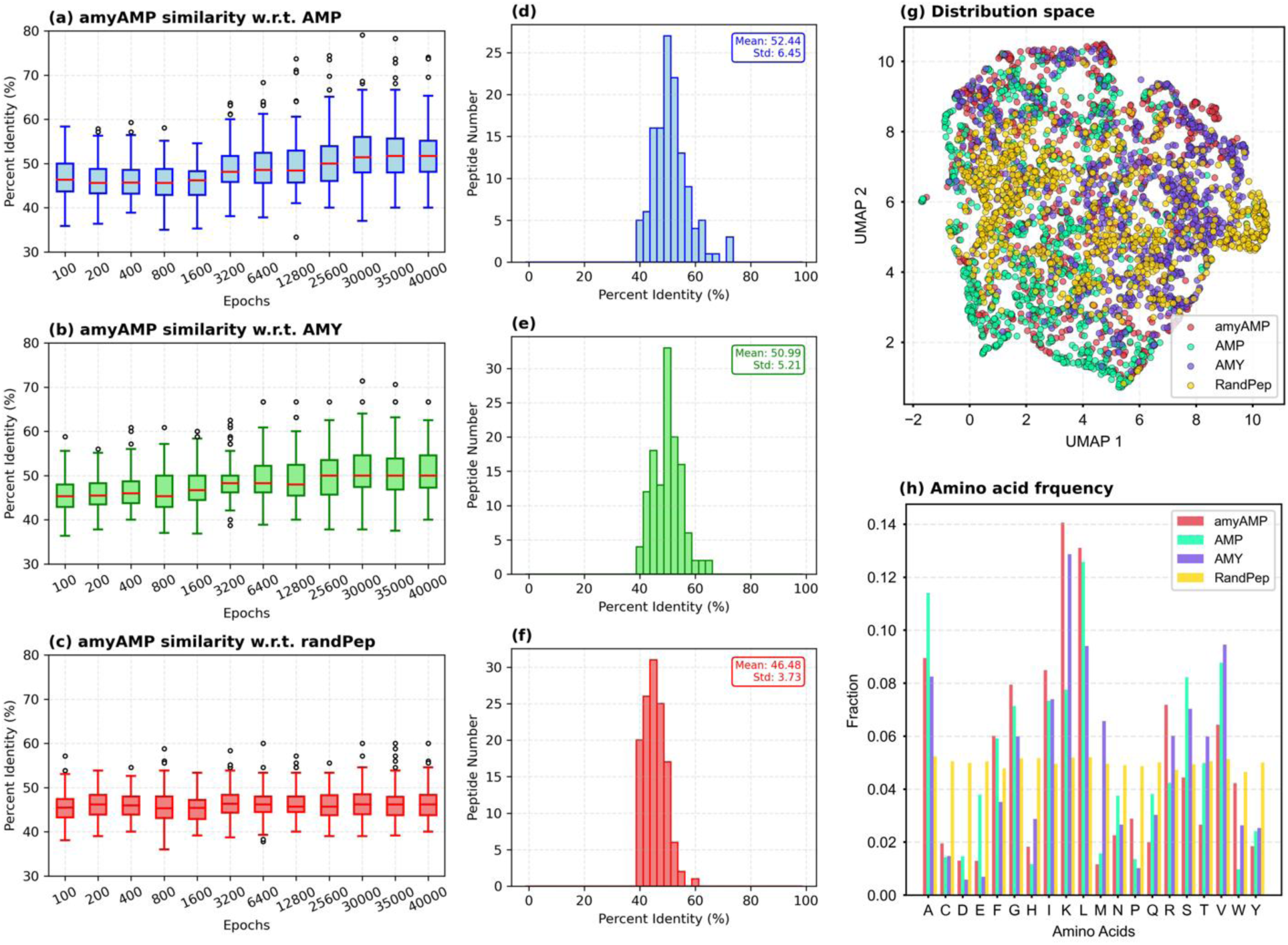
Model learning statistics. The percentage identities of generated peptides with respect to training datasets are shown at different training epochs. (a) antimicrobial peptides (AMPs), (b) amyloid peptides (AMYs), and (c) random peptides (RandPep). Panels (d–f) illustrate the distribution of percent identities between generated peptides and the training datasets at the final epochs. The generated sequences exhibit substantial similarity to AMPs (52.4%) and AMYs (51%), whereas similarity to random peptides remains low (46.4). This enrichment suggests that the generated peptides inherit both antimicrobial and amyloidogenic characteristics. Panel (g) shows the distribution of different peptide groups, highlighting the overlap of amyAMP sequences that incorporate both AMP and amyloid properties. Panel (h) compares the amino acid frequency distributions of generated and training datasets.

Upon completion of training, the generator component of the amyAMP model was used to produce 1000 novel peptide sequences. To assess whether the model had effectively learned the sequence-property relationships encoded in the training data, we performed dimensionality reduction using Uniform Manifold Approximation and Projection (UMAP) and compared the distribution of generated peptides with those of the training and reference datasets (Figure 2g). The resulting embedding revealed that amyAMP-generated peptides (red) occupied a region of the latent space that substantially overlapped with both the AMP and AMY training datasets. In contrast, randomly generated peptides formed a distinct cluster, indicating that the amyAMP-generated sequences reside within the biologically relevant peptide space rather than the random sequence landscape. To further validate this observation, the embedding analysis was repeated by combining the AMP and AMY datasets into a unified reference distribution (Figure S2), which yielded a consistent overlap with the amyAMP-generated peptides. Together, these results indicate that the amyAMP model effectively captures the latent structural and physicochemical features characteristic of peptides possessing both antimicrobial and amyloidogenic properties.

Consistent with the latent space analysis, amino acid frequency profiling (Figure 2h) showed that the residue composition of amyAMP-generated peptides closely resembles that of the training datasets. Preserving these residue- and peptide-level statistics provides additional evidence that the model effectively captured the underlying sequence patterns in the training data while generating novel sequences. At the same time, variability observed among the generated peptides suggests that the model explores a diverse region of the learned sequence space rather than simply reproducing sequences from the training datasets.

### 3.2 Model assessment on amyloid-antimicrobial relative properties

To evaluate whether amyAMP-generated peptides reproduce the defining biophysical signatures of antimicrobial and amyloidogenic sequences, we quantified and compared nine key related physicochemical properties across all four peptide datasets: amyAMP-generated peptides, AMPs, AMYs and RandPep. As shown in Figure 3, amyAMP peptides exhibit elevated net charge, AMP-favorable residue composition, high isoelectric points, and increased hydrophobic moment, closely aligning with the natural AMP distribution. Concurrently, amyAMP sequences display enhanced β-sheet propensity, amphipathicity, and GRAVY hydrophobicity, features commonly associated with amyloidogenic peptide assembly. Shannon entropy and instability index values further show that amyAMP peptides maintain structural and compositional diversity, distinguishing them from both natural reference sets. Collectively, these multidimensional property profiles demonstrate that amyAMP effectively learns and reproduces the dual physicochemical landscape underlying antimicrobial function and amyloid formation, validating the biological plausibility of the generated peptide library. Furthermore, we analyzed the correlations among multiple AMP- and AMY-related physicochemical properties shown in Figure S3. The amyAMP-generated peptides exhibited correlation patterns highly consistent with those observed in both the AMP and AMY datasets, whereas random peptides showed weak or inconsistent correlations across the same properties. These findings indicate that the model not only learns individual property distributions but also captures the underlying inter-property relationships characteristic of bioactive antimicrobial and amyloidogenic peptides.

**Figure 3.**
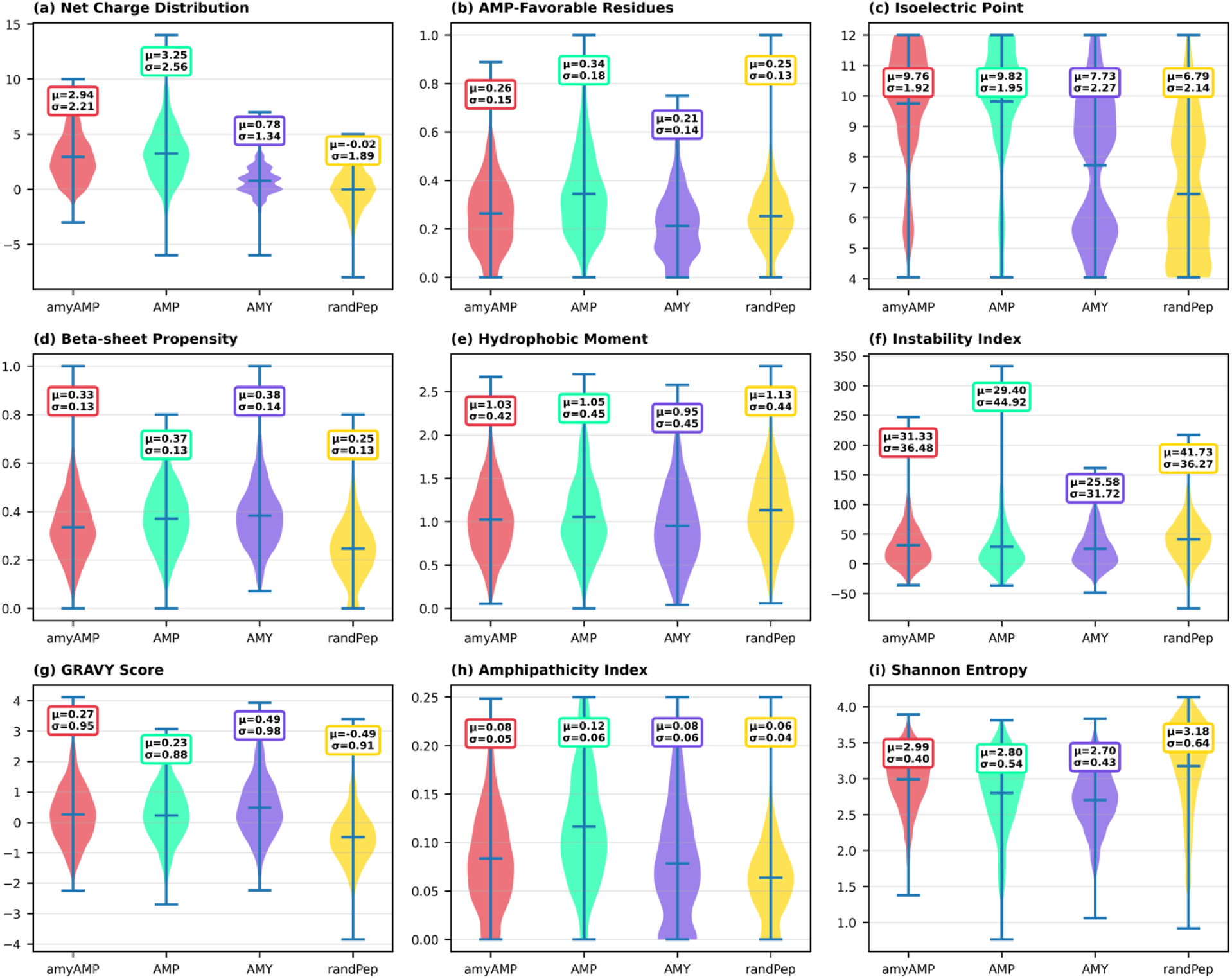
Comparative physicochemical property profiles of amyAMP-generated peptides versus reference peptide sets. Violin plots summarize nine antimicrobial- and amyloid-relevant physicochemical properties across four datasets: amyAMP-generated peptides, antimicrobial peptides (AMPs), amyloidogenic peptides (AMYs), dataset used in training and randomly peptides (RandPeps). Properties evaluated include: (a) Net charge, (b) Fraction of AMP-favorable residues, (c) Isoelectric point, (d) β-sheet propensity, (e) Hydrophobic moment, (f) Instability index, (g) GRAVY hydrophobicity score, (h) Amphipathicity index, and (i) Shannon entropy. The distributional patterns reveal that amyAMP-generated peptides share key physicochemical signatures with both AMP and amyloid datasets, such as elevated charge, β-sheet propensity, hydrophobic moment, and amphipathicity while remaining distinct from random peptides. These analyses confirm that the amyAMPmodel successfully captures dual antimicrobial and amyloidogenic physicochemical trends characteristic of functional bioactive peptides.

In addition, we examined the distributions of the six physicochemical features comprising the PC6 encoding. As shown in Figure S4, amyAMP-generated peptides closely match the statistical profiles of the reference AMP and AMY datasets, mirroring the trends observed in the broader antimicrobial- and amyloid-related property analyses. The generated peptides consistently reproduce the balanced property distributions of the training data, indicating that the model captures underlying physicochemical patterns rather than memorizing individual sequences. In contrast, random peptides show markedly different and often extreme distributions across all six properties, underscoring the specificity learned by amyAMP.

### 3.3 In-silico validation of dual-functional peptides generated by amyAMP

To further assess the functional credibility of amyAMP-generated peptides, we evaluated their predicted activities using four independent deep-learning based predictors trained on antimicrobial or amyloidogenic peptide datasets. Such external predictors provide an unbiased computational framework to determine whether the generated sequences capture the sequence and physicochemical features characteristic of functional antimicrobial peptides and amyloid-forming peptides. Specifically, antimicrobial activity was assessed using AI4AMP and AMPScanner, while amyloidogenic propensity was evaluated using Waltz and AmyloGram.^45,50,51,60,61^

Across all external models, amyAMP-generated peptides exhibited strong and consistent enrichment in high-activity prediction categories (Figure 4 and Figure S5). Both AI4AMP and AMPScanner, widely used AMP classifiers, assigned the majority of generated sequences to the high (0.7–0.9) and very high (0.9–1.0) antimicrobial probability classes, suggesting that the model effectively captures sequence patterns associated with antimicrobial activity. Complementarily, Waltz and AmyloGram classified a substantial fraction of amyAMP peptides as possessing high or very high amyloidogenic propensity. The concordance across these independent predictors indicates that the dual-functional design objective, simultaneous antimicrobial and amyloid-forming potential, was preserved in the generated peptide library.

**Figure 4.**
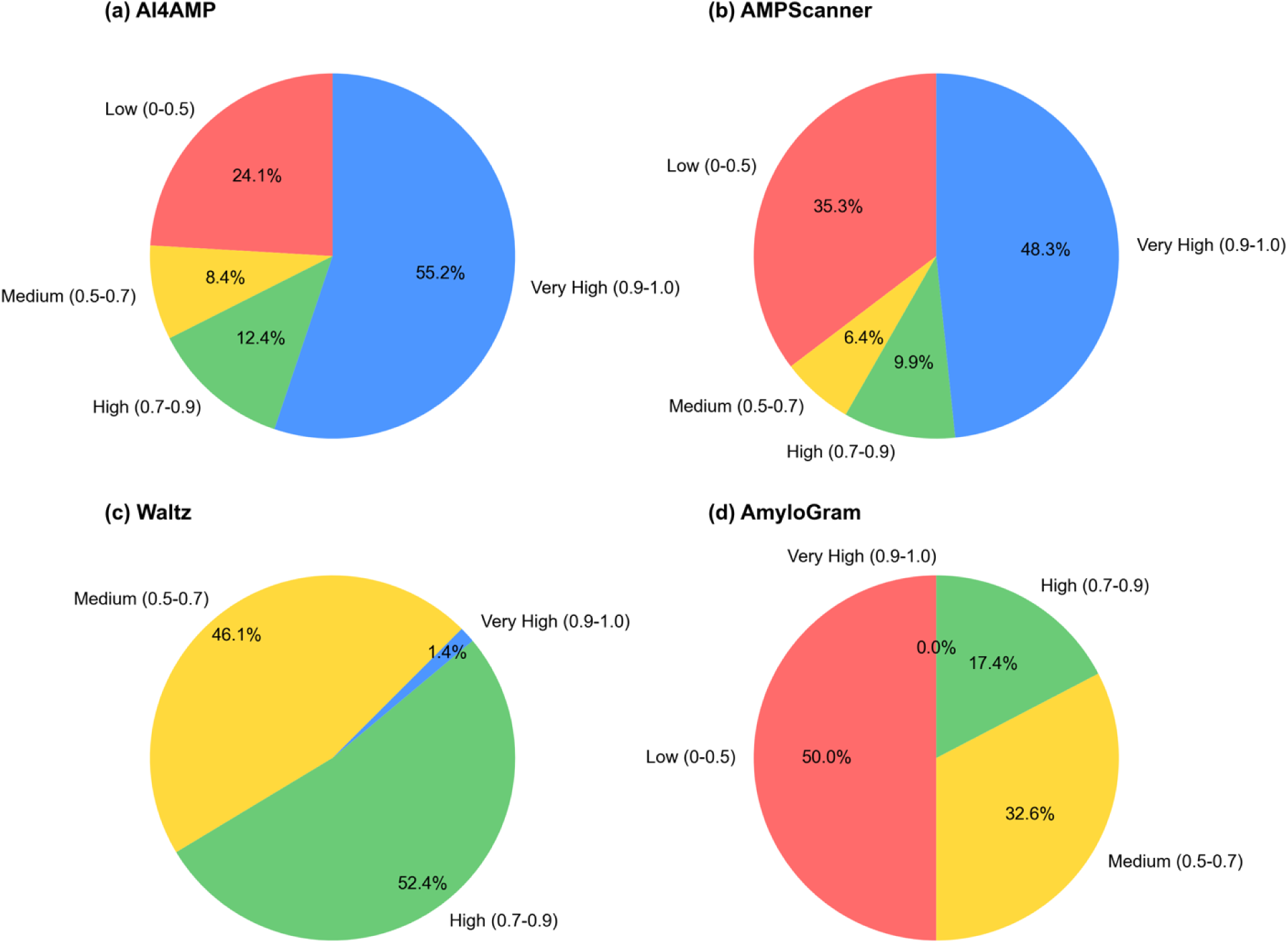
Independent computational validation of amyAMP-generated peptides using multiple deep-learning predictors. Predicted activity scores were grouped into four categories: low (0–0.5), medium (0.5–0.7), high (0.7–0.9), and very high (0.9–1.0). Each panel presents a pie chart showing the percentage distribution of amyAMP-generated peptides across these score categories. (a) Predictions from AI4AMP indicate that most sequences fall within the high and very high antimicrobial activity categories. (b) AMPScanner shows a similar enrichment of peptides with high antimicrobial scores. (c) Amyloid propensity predicted by Waltz reveals a strong shift toward high and very high amyloidogenic probabilities with no low-scored peptides. (d) Predictions from AmyloGram likewise classify the majority of peptides within the medium or high amyloidogenic categories.

Notably, the enrichment of high-scoring peptides across both antimicrobial and amyloid predictors suggests that the amyAMP model has learned shared physicochemical and structural determinants underlying these functional classes, including amphipathicity, β-sheet–promoting sequence patterns, hydrophobic moment, and charge distribution. These properties are known to play key roles in both antimicrobial membrane interactions and amyloid fibril formation. Collectively, these results provide strong computational evidence that amyAMP-generated sequences possess integrated bioactivity signatures, supporting their prioritization for downstream experimental validation.

### 3.4 Molecular dynamics simulations

While statistical analyses and external deep-learning predictors suggested strong dual antimicrobial and amyloidogenic potential of amyAMP-generated peptides, we further assessed their functional behavior using physics-based molecular dynamics simulations. Specifically, coarse-grained simulations were performed to investigate peptide-membrane interactions and peptide self-assembly, enabling direct comparison with characterized antimicrobial and amyloid peptides from the training datasets. From the library of 1000 amyAMP-generated sequences, the top 100 peptides were used for simulation. For benchmarking, a reference set comprising 50 peptides from each of three experimentally defined activity classes, low, intermediate, and high MIC, was extracted from the training dataset, facilitating direct comparison with experimentally validated antimicrobial peptides.

#### (i) Peptide-membrane binding simulations

Each amyAMP-generated peptide and reference peptide was simulated in its monomeric form for 1 µs in the presence of a Gram-positive bacterial model lipid bilayer composition. At the start of each simulation, peptides were positioned approximately 20 Å above the membrane surface in the aqueous phase. Across nearly all simulations, peptides rapidly diffused toward the membrane and adsorbed onto the lipid bilayer within the first few nanoseconds. Once bound, the peptides remained stably associated with the membrane throughout the remainder of the trajectories. Representative snapshots illustrating the binding process are shown in Figure 5 (a–c). Peptides initially located in the solvent phase reoriented upon approaching the membrane and subsequently adopted conformations in which their hydrophobic faces were oriented toward the lipid core while polar residues remained exposed to the lipid headgroup region.

**Figure 5.**
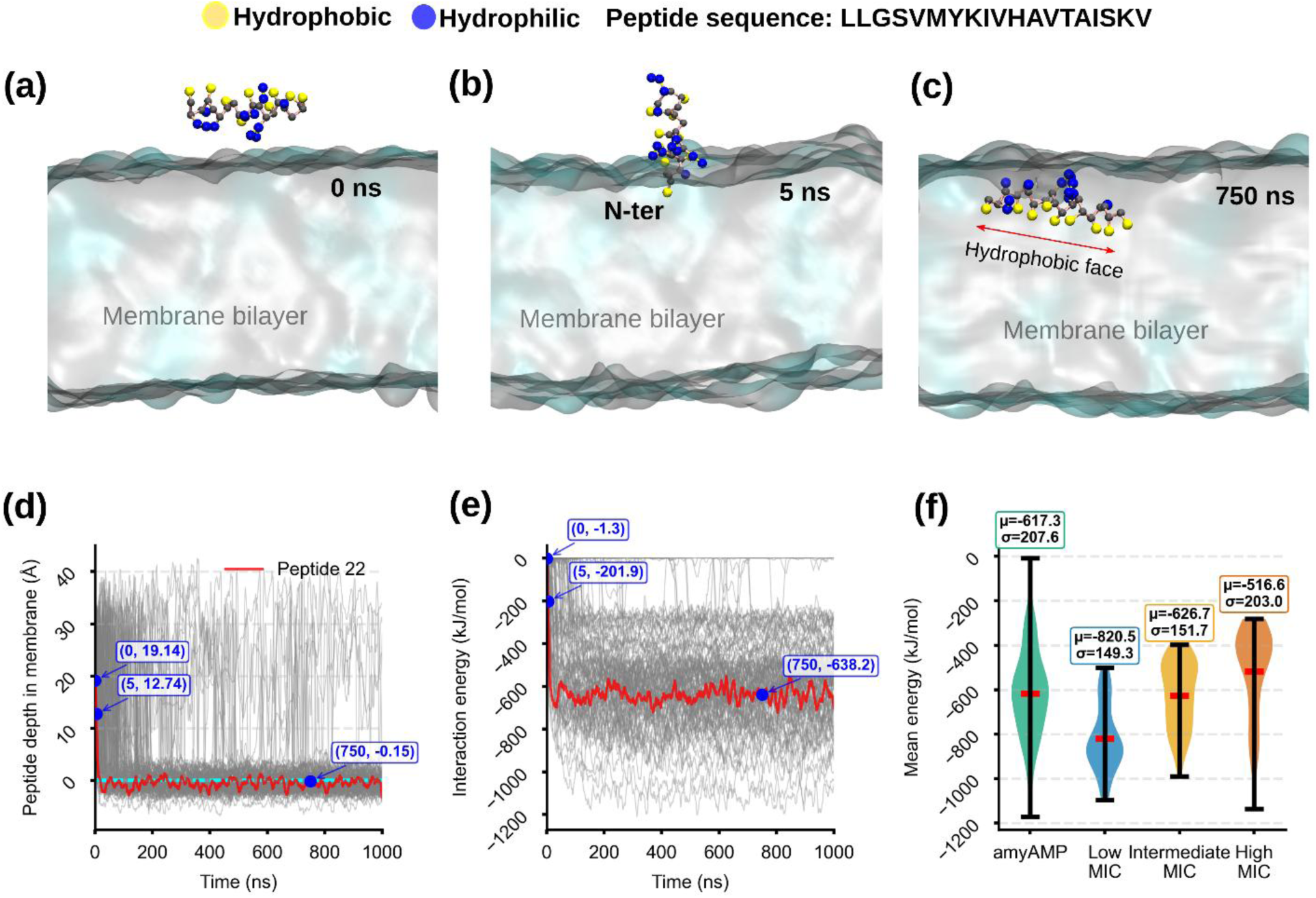
Peptide-membrane interaction dynamics from coarse-grained molecular dynamics simulations. Representative snapshots, captured at (a) 0 ns (b) 5 ns and (c) 750 ns, amyAMP-generated single peptide simulation illustrating rapid adsorption from the aqueous phase followed by partial insertion into the lipid bilayer. (d) Insertion depth profiles of amyAMP-generated peptides relative to the membrane surface, showing that most peptides are stabilized at a shallow depths (∼3–5 Å below the bilayer interface). (e) Time evolution of peptide–lipid interaction energy during the simulation. The red trace in panels (d–e) corresponds to the trajectory from which the snapshots in (a–c) were extracted. (f) Distribution of mean peptide-lipid interaction energies for amyAMP-generated peptides compared with experimentally characterized antimicrobial peptides grouped by minimum inhibitory concentration (MIC), demonstrating overlapping membrane-binding energetics across activity classes.

In addition to qualitative observations of membrane binding, peptide insertion behavior was quantified across all simulations. A representative trajectory, from which snapshots were extracted, is shown as an example. Analysis of peptide depth relative to the bilayer surface revealed that most amyAMP-generated peptides rapidly adsorbed and partially inserted into the membrane medium. As shown in Figure 5d, peptides predominantly occupied shallow insertion depths of approximately 3–5 Å below the membrane surface rather than penetrating deeply into the hydrophobic core. This shallow insertion pattern is consistent with the amphipathic nature of antimicrobial peptides, in which hydrophobic residues interact with the lipid core while polar or charged residues stabilize interactions with the membrane headgroups.

To quantify the strength of peptide-membrane interactions, peptide-lipid interaction energies were computed throughout the trajectories. Across the 100 amyAMP-generated peptides, interaction energies exhibited substantial sequence-dependent variability, ranging from approximately −200 to −1200 kJ mol⁻¹, with a mean value near −600 kJ mol⁻¹ (Figure 5(e–f)). Importantly, the energy distribution of amyAMP-generated peptides overlapped with that observed for experimentally characterized antimicrobial peptides spanning the low-, intermediate-, and high-MIC categories. This overlap suggests that the generated sequences reproduce a broad range of membrane-binding strengths associated with experimentally validated antimicrobial activity.

For each simulations, the mean peptide-lipid interaction energy, calculated as the sum of van der Waals and electrostatic contributions, was averaged over the final 200 ns of simulation to ensure equilibration. Comparison of these mean energies revealed that peptides associated with low MIC values displayed more favorable (i.e., more negative) interaction energies and a higher frequency of strong membrane-binding events relative to intermediate- and high-MIC peptides. Notably, the amyAMP-generated peptides spanned the entire energetic range observed for the reference peptides (Figure 5(f)), with more than half of the generated sequences exhibiting interaction energies comparable to those of low- and intermediate-MIC antimicrobial peptides. The trajectories of peptide depth in membrane for refernces sets are shown in figure S6. Together, these results indicate that amyAMP-generated peptides occupy a membrane-binding regime consistent with experimentally validated antimicrobial peptides, supporting the functional relevance of the generated sequence library.

#### (ii) Peptide self-assembly simulations

To evaluate the intrinsic aggregation propensity of amyAMP-generated peptides, coarse-grained molecular dynamics simulations were performed to probe peptide self-assembly behavior in solution. For each of the 100 amyAMP sequences, eight identical copies of the peptide were randomly placed in an aqueous simulation box and simulated for 1 µs per system. At the beginning of the simulations, all peptides were randomly dispersed in the simulation box. However, peptide-peptide interactions rapidly emerged, leading to the formation of oligomeric assemblies within the first ∼100 ns of the trajectories (Figure 6). Representative snapshots from one simulation are shown in Figure 6a–c. The peptide selected to illustrate membrane insertion in the context of antimicrobial activity was also used here to demonstrate aggregation and self-assembly behavior. Peptides initially dispersed in solution quickly initiated intermolecular contacts, forming small oligomeric clusters within ∼60 ns (Figure 6b), which subsequently evolved into compact assemblies characterized by a hydrophobic core (Figure 6c). Such structural organization is consistent with the self-association patterns commonly observed in amyloid-forming peptides.

**Figure 6.**
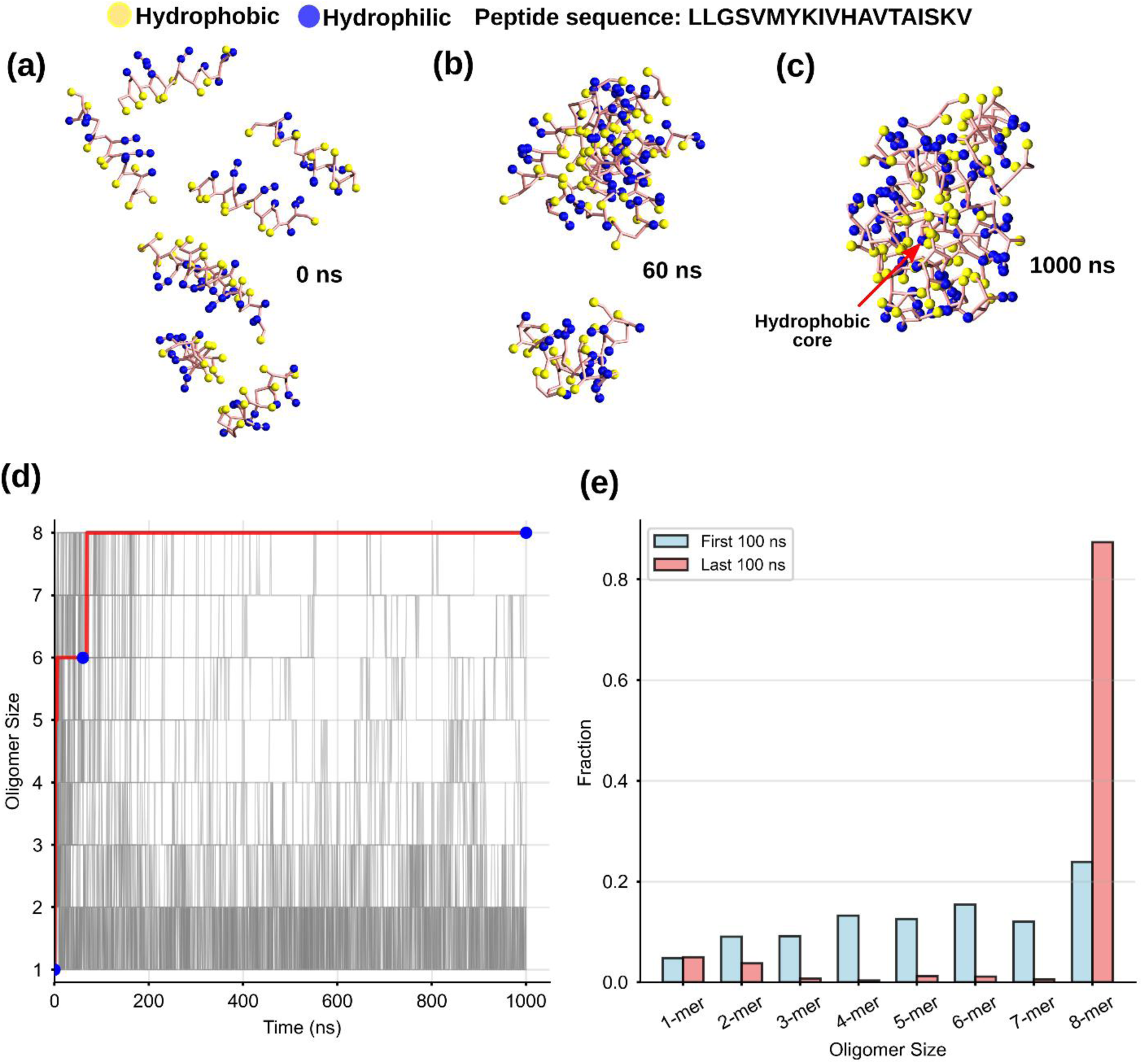
Aggregation behavior of amyAMP-generated peptides. Representative snapshots, captured at (a) 0 ns (b) 60 ns (c) 1000 ns, showing rapid self-assembly of eight peptides from dispersed monomeric state (0 ns) to compact aggregates with a hydrophobic core (1000 ns). (d) Time evolution of oligomer size during the 1µs simulation, indicating fast formation of higher-order assemblies. (e) Oligomer size distribution comparing the first and last 100 ns, showing that >80% of peptides form stable 8-mer aggregates in the final stage, consistent with strong amyloidogenic propensity.

To quantitatively characterize the self-assembly process, oligomerization states were monitored throughout the simulation trajectories. Analysis of oligomer populations revealed progressive self-assembly, with most systems converging toward the formation of large oligomeric clusters approaching where eight peptides had assembled during the final 100 ns of simulation (Figure 6(d-e)). In a minority of trajectories, individual peptides exhibited delayed incorporation into the aggregate but ultimately joined the largest dominant stable oligomer. Overall, these simulations demonstrate a strong intrinsic self-assembly trends among amyAMP-generated peptides, supporting the predicted amyloidogenic character of the designed sequences.

#### (iii) Membrane disruption simulations

To further investigate whether amyAMP-generated peptides simultaneously exhibit antimicrobial membrane activity and aggregation behavior, membrane disruption simulations were performed under conditions mimicking elevated peptide concentration. Analysis of peptide-membrane binding simulations showed that the interaction energies of amyAMP-generated peptides span the full range observed for peptides associated with low-, intermediate-, and high-MIC classes, with a clear dependence on peptide length. To probe dual functionality across this broad interaction-energy landscape, three representative peptides were selected from distinct regions of the interaction energy distribution while also capturing sequence-length diversity. These peptides were designated as low-, intermediate-, and high-MIC analogs. Each system comprised nine identical peptide molecules initially positioned in the aqueous phase above the membrane surface and simulated for 10 µs using coarse-grained molecular dynamics. Simulations were carried out using both Gram-positive and Gram-negative membrane models to assess the influence of membrane composition on peptide activity.

Consistent with the behavior observed in the peptide-membrane binding simulations, peptides rapidly diffused toward the membrane surface and adsorbed onto the lipid bilayer during the early stages of the trajectories (Figure 7). Following adsorption, peptides established intermolecular contacts and progressively formed oligomeric clusters on the membrane surface. However, the extent of clustering and the resulting membrane perturbation varied markedly among the three peptides. The high-MIC analog exhibited relatively weak peptide-peptide interactions and failed to assemble into large surface-associated clusters (Figure 7a). Consequently, it induced minimal membrane disruption, with only modest changes in membrane thickness and curvature (Figure 7b–c).

**Figure 7.**
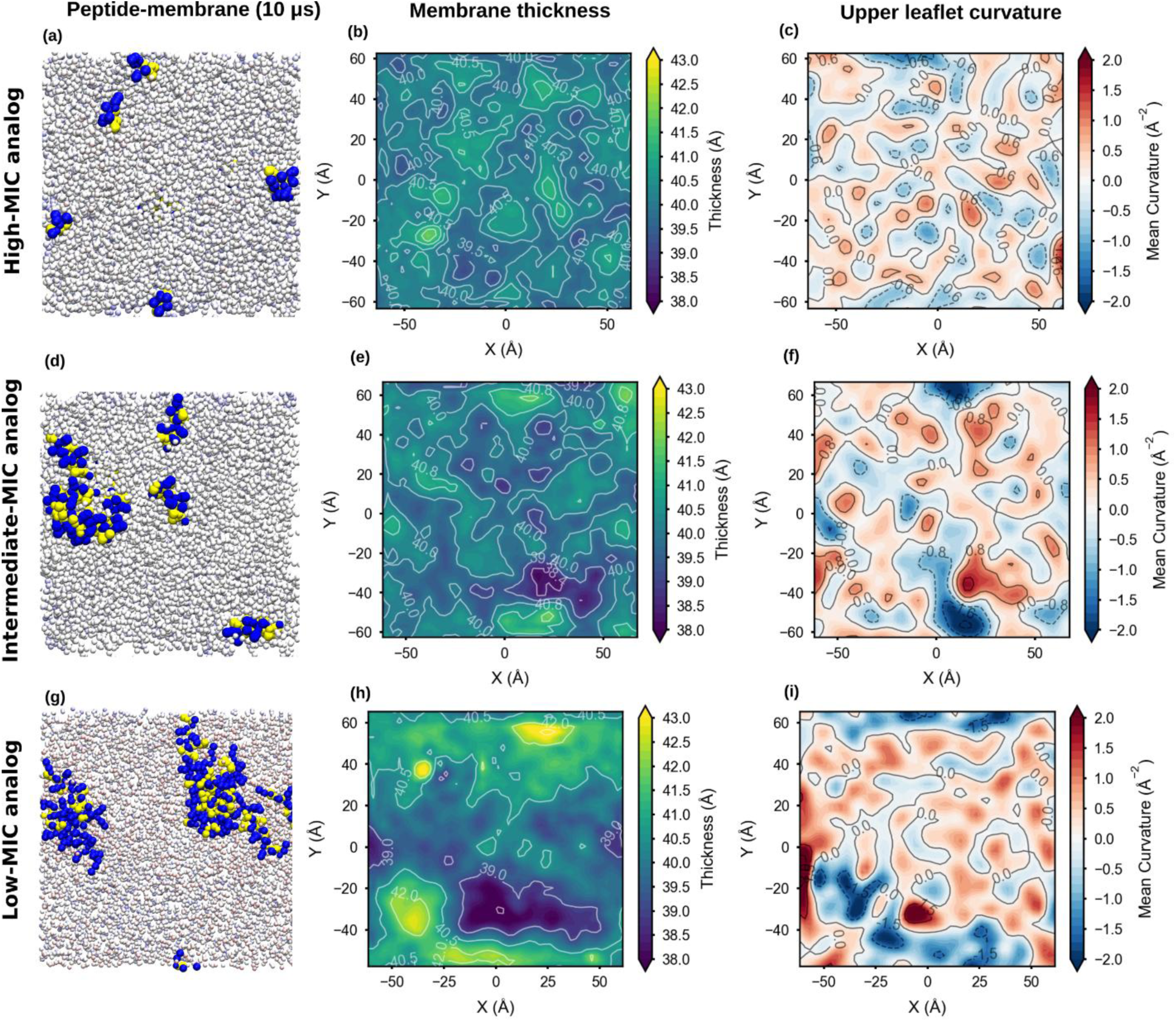
Membrane disruption induced by amyAMP peptides of different sequence lengths. Three selected amyAMP-generated peptides, high-, intermediate-, and low-MIC analogs, were simulated with nine identical copies of peptides initially positioned above the Gram-positive membrane surface. The high-MIC peptide (a–c) forms only weak peptide-peptide interactions on the membrane surface and does not assemble into large oligomeric clusters, resulting in minimal membrane perturbation. In contrast, the low-MIC peptide (g–i) forms a large surface-associated peptide cluster that induces pronounced membrane thinning and strong curvature, indicating substantial disruption of membrane integrity. The intermediate-MIC peptide (d–f) displays intermediate behavior between these two extremes. Together, these results demonstrate that cooperative peptide self-assembly on the membrane surface plays a critical role in determining the extent of membrane disruption.

In contrast, the intermediate- and low-MIC analogs formed substantially larger oligomeric assemblies on the membrane surface (Figure 7d–i). In these systems, peptides organized into clusters with extended hydrophobic interfaces that engaged strongly with the lipid bilayer. Such cooperative assemblies induced pronounced membrane perturbations, including localized thinning and significant changes in mean curvature. The disruptive effect was most pronounced for the low-MIC analog (Figure 7g–i), which formed the largest surface-associated cluster and exhibited the strongest membrane deformation. These results indicate that strong peptide-membrane interactions alone are insufficient to drive membrane disruption; rather, effective disruption requires the combined effect of strong membrane binding and cooperative peptide clustering.

The primary results shown in the main text correspond to the Gram-positive membrane model (Figure 7), while analogous simulations with Gram-negative membrane mimetics are presented in the Supplementary Information (Figure S7). Compared with the Gram-positive system, peptide adsorption and clustering were reduced on the Gram-negative membrane, leading to weaker membrane thinning and smaller curvature changes. This diminished disruption likely reflects differences in lipid composition and electrostatic interactions that modulate peptide binding affinity. Collectively, these findings demonstrate that amyAMP-generated peptides can simultaneously exhibit aggregation-driven self-assembly and membrane-active antimicrobial behavior, and further establish that cooperative clustering in conjunction with strong peptide-membrane interactions is a key determinant of membrane disruption, supporting the dual-functional design principle of the amyAMP model.

To quantify the prevalence of dual functionality within the amyAMP-generated library, peptides were classified based on two mechanistic criteria: peptide-membrane interaction energy (< −600 kJ mol⁻¹) and aggregation propensity (aggregate size ≥ 5 during the final 0.2 µs of simulation). As shown in Figure 8a, a substantial fraction of peptides satisfies both conditions, yielding a subset of 51 dual-functional sequences that combine strong membrane affinity with robust self-assembly behavior. Comparison of amino acid composition between the full peptide set and the dual-functional subset (Figure 8b) revealed highly similar residue distributions, with enrichment of hydrophobic and positively charged residues in the dual-functional peptides. These features are consistent with enhanced membrane interaction and aggregation propensity. Together, these results demonstrate that dual functionality is not restricted to a small subset of sequences but is broadly represented within the generated library, indicating that the amyAMP model effectively captures the sequence determinants underlying both antimicrobial activity and amyloid-like self-assembly.

**Figure 8.**
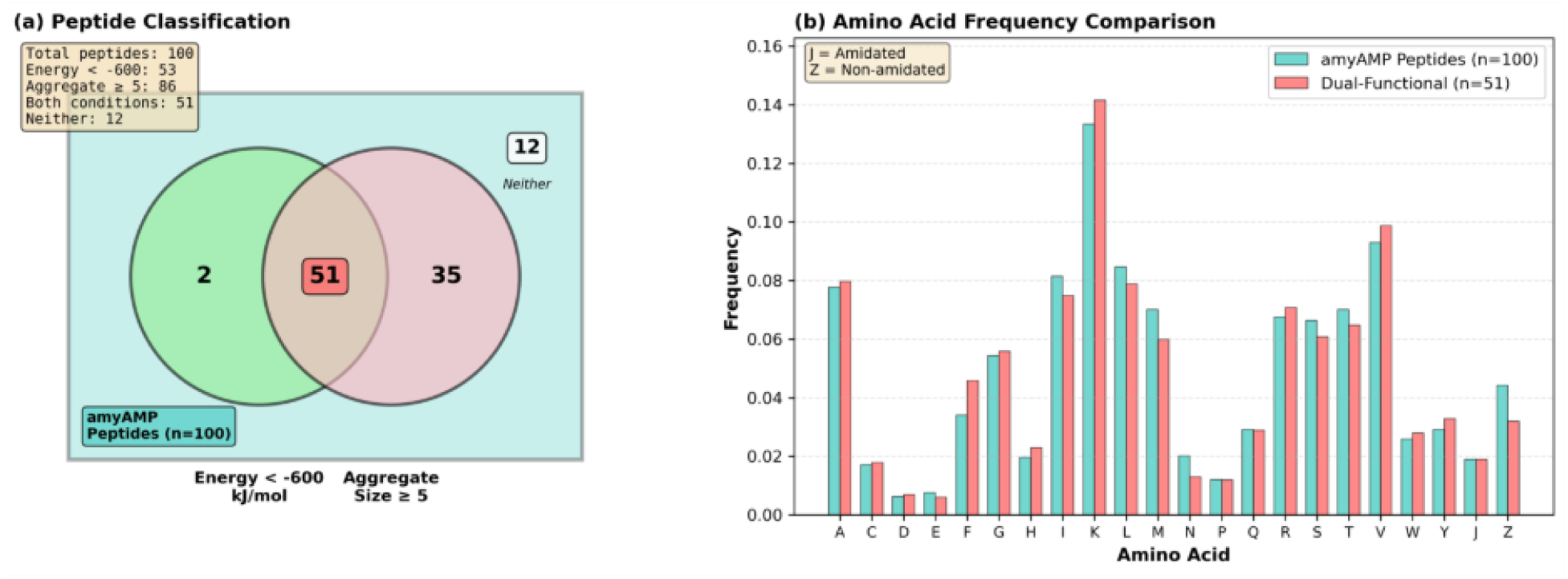
Classification and amino acid composition of amyAMP peptides based on aggregation and membrane interaction. (a) Venn diagram of 100 amyAMP peptides classified by strong membrane interaction energy (< −600 kJ mol⁻¹) and high aggregation propensity (aggregate size ≥ 5). The overlap (red, n = 51) represents dual-functional peptides exhibiting both properties. (b) The distribution of amino acid composition of all peptides (teal, n = 100) compared with the dual-functional subset (red, n = 51) are shown. Dual-functional peptides are enriched in hydrophobic and positively charged residues, consistent with enhanced membrane interaction and aggregation.

## 5. Conclusion and Future Perspectives

The emergence of antimicrobial resistance has created an urgent need for alternative therapeutic strategies beyond conventional small-molecule antibiotics, with antimicrobial peptides representing one of the most promising classes of next-generation therapeutics. At the same time, increasing evidence suggests a fundamental and underexplored connection between antimicrobial activity and amyloid-like self-assembly, where shared physicochemical features, such as amphipathicity, β-sheet propensity, and charge distribution, govern both membrane activity and aggregation behavior.^15–17^ However, the rational design of peptides that simultaneously encode these dual functionalities remains a major challenge due to the complexity of the underlying sequence-structure-function relationships.

In this work, we introduced amyAMP, a bidirectional generative adversarial network that enables the de novo design of peptides integrating antimicrobial activity with amyloidogenic self-assembly. By jointly learning from antimicrobial and amyloid peptide datasets, the model captures the latent biochemical landscape governing dual functionality. Latent-space analysis demonstrated that amyAMP-generated peptides occupy a sequence-property space that closely overlaps with biologically relevant peptide datasets while remaining distinct from random sequences (Figure 2g), indicating successful learning of functionally meaningful representations. Multi-level validation further supports the functional relevance of the generated peptides. Independent deep-learning predictors consistently classified amyAMP sequences within high antimicrobial and amyloidogenic propensity regimes (Figure 4), highlighting the robustness of the learned features across orthogonal models. Importantly, coarse-grained molecular dynamics simulations provided physics-based mechanistic validation. The simulations revealed rapid membrane adsorption, stable amphipathic insertion, and strong peptide-lipid interactions comparable to experimentally validated AMPs (Figure 5). Concurrently, peptide self-assembly simulations demonstrated robust oligomerization and the formation of compact aggregates (Figure 6), consistent with amyloid-like behavior. Notably, membrane disruption simulations established that effective antimicrobial activity emerges from the cooperative interplay between peptide-membrane affinity and peptide-peptide aggregation, where the formation of large surface-associated clusters drives membrane thinning and curvature perturbations (Figure 7). These findings provide mechanistic evidence that antimicrobial function and amyloid-like self-assembly are not independent phenomena but are intrinsically coupled through shared physicochemical determinants.

Beyond methodological advancement, this work establishes a unified framework for multifunctional peptide design by integrating generative artificial intelligence with physics-based simulations, thereby bridging sequence-level learning and mechanistic insight. This approach addresses a key limitation of existing strategies that rely on either data-driven prediction or simulation-based validation in isolation. By combining these paradigms, amyAMP enables efficient exploration of complex sequence-function landscapes that are otherwise difficult to access through conventional methods. Future efforts will focus on experimental validation of the designed peptides to confirm antimicrobial activity, aggregation behavior, and structural properties. Further improvements may be achieved by incorporating structure-aware generative models, protein language models, and simulation-guided optimization frameworks. Expanding the design space to include toxicity, stability, and functional specificity will be critical for therapeutic translation. Overall, this integrative strategy provides a scalable platform for the discovery of multifunctional biomolecules and offers a promising direction for developing next-generation peptide therapeutics to combat antimicrobial resistance.

## Acknowledgements

This work was supported by the Monash Data Futures Institute (MDFI) Seed Grant, Monash University, and the Institute Postdoctoral Fellowship at IIT Bombay. The authors thank Dr. Senthil Arumugam, a member of EMBL Australia, Monash University and Prof. Andrea Collevecchio, School of Mathematics, Monash University for their assistance during the early stages of the project.

## Code availability

The source code for the amyAMP model and all associated analysis is publicly available at: https://github.com/anupkprasad/amyAMP

## Data availability

All peptide sequences used in this study, including training datasets and amyAMP-generated peptides, are available at: https://github.com/anupkprasad/amyAMP.

