## Supplementary material for "Generative Deep Learning and Molecular Dynamics Reveal Design Principles for Amyloid-Like Antimicrobial Peptides"

Current address:

**Table S1: Physicochemical Properties of Amino Acids (H1–NCI)**

(H1: hydrophobicity, V: side-chain volume, P1: polarity, Pl: pH at isoelectric point, pKa: dissociation constant of –COOH group, NCI: net charge index), J: amdiation, Z: non-amidation

| Amino Acid | H1 (Hydrophobicity) | V (Side-chain Volume) | P1 (Polarity) | Pl (pI) | pKa (COOH) | NCI (Net Charge Index) |
|---|---|---|---|---|---|---|
| A | 0.62 | 27.5 | 8.1 | 6.00 | 2.34 | 0.007187 |
| C | 0.29 | 44.6 | 5.5 | 5.07 | 1.96 | -0.03661 |
| D | -0.90 | 40.0 | 13.0 | 2.77 | 1.88 | -0.02382 |
| E | -0.74 | 62.0 | 12.3 | 3.22 | 2.19 | 0.006802 |
| F | 1.19 | 115.5 | 5.2 | 5.48 | 1.83 | 0.037552 |
| G | 0.48 | 0.0 | 9.0 | 5.97 | 2.34 | 0.179052 |
| H | -0.40 | 79.0 | 10.4 | 7.59 | 1.82 | -0.01069 |
| I | 1.38 | 93.5 | 5.2 | 6.02 | 2.36 | 0.021631 |
| K | -1.50 | 100.0 | 11.3 | 9.74 | 2.18 | 0.017708 |
| L | 1.06 | 93.5 | 4.9 | 5.98 | 2.36 | 0.051672 |
| M | 0.64 | 94.1 | 5.7 | 5.74 | 2.28 | 0.002683 |
| N | -0.78 | 58.7 | 11.6 | 5.41 | 2.02 | 0.005392 |
| P | 0.12 | 41.9 | 8.0 | 6.30 | 1.99 | 0.239531 |
| Q | -0.85 | 80.7 | 10.5 | 5.65 | 2.17 | 0.049211 |
| R | -2.53 | 105.0 | 10.5 | 10.76 | 2.17 | 0.043587 |
| S | -0.18 | 29.3 | 9.2 | 5.68 | 2.21 | 0.004627 |
| T | -0.05 | 51.3 | 8.6 | 5.60 | 2.09 | 0.003352 |
| V | 1.08 | 71.5 | 5.9 | 5.96 | 2.32 | 0.057004 |
| W | 0.81 | 145.5 | 5.4 | 5.89 | 2.83 | 0.037977 |
| Y | 0.26 | 117.3 | 6.2 | 5.66 | 2.20 | 0.023599 |
| J | 1.00 | 1.0 | 1.0 | 1.00 | 1.00 | 1.000000 |
| Z | 2.00 | 2.0 | 2.0 | 2.00 | 2.00 | 2.000000 |

### S1. Two modular neural building blocks were employed to construct the amyAMP architecture

1. **TBR (Transposed Block with Batch Normalization and ReLU)** consisting of a two-dimensional transposed convolution layer (*ConvTranspose2D*), followed by batch normalization and a ReLU activation function. This block facilitates upsampling and feature refinement in the Generator.
2. **CR (Convolutional Block with LeakyReLU)** composed of a two-dimensional convolution layer (*Conv2D*) coupled with a LeakyReLU activation function, enabling hierarchical feature extraction in the Encoder and Discriminator.

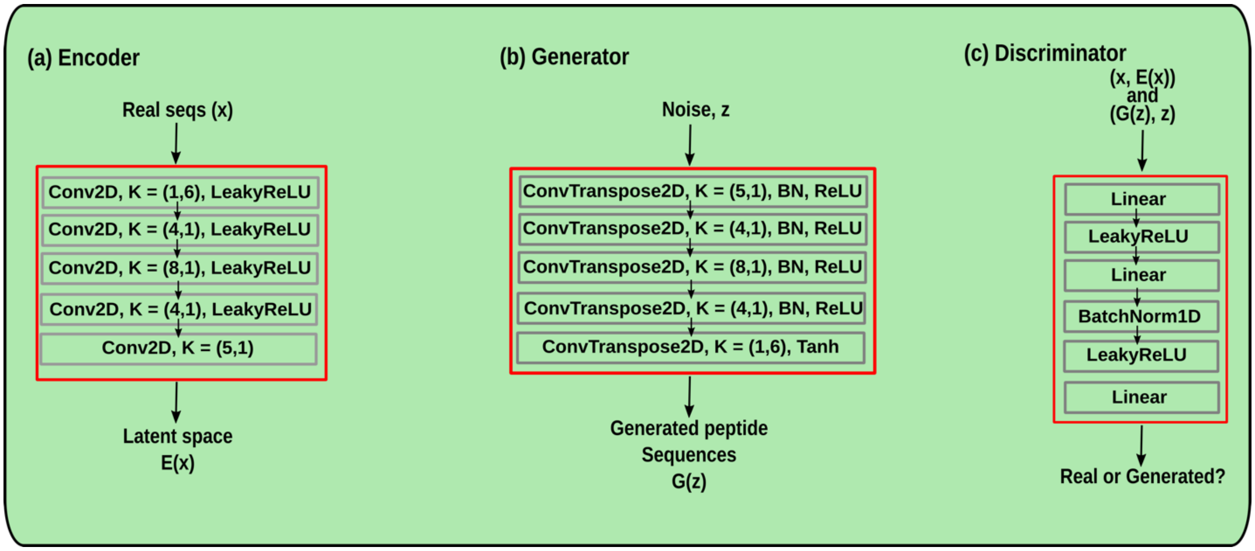


**Figure S1. Architecture of the amyAMP machine-learning model.** (a) Encoder: composed of 2D convolutional layers with LeakyReLU activation and varying kernel sizes for hierarchical feature extraction. (b) Generator: built using transposed convolutional layers with ReLU activation and progressively increasing kernel sizes to reconstruct peptide feature maps from latent vectors. (c) Discriminator: implemented as a series of linear layers with LeakyReLU activation and batch normalisation, enabling effective discrimination between real and generated peptide–latent pairs.

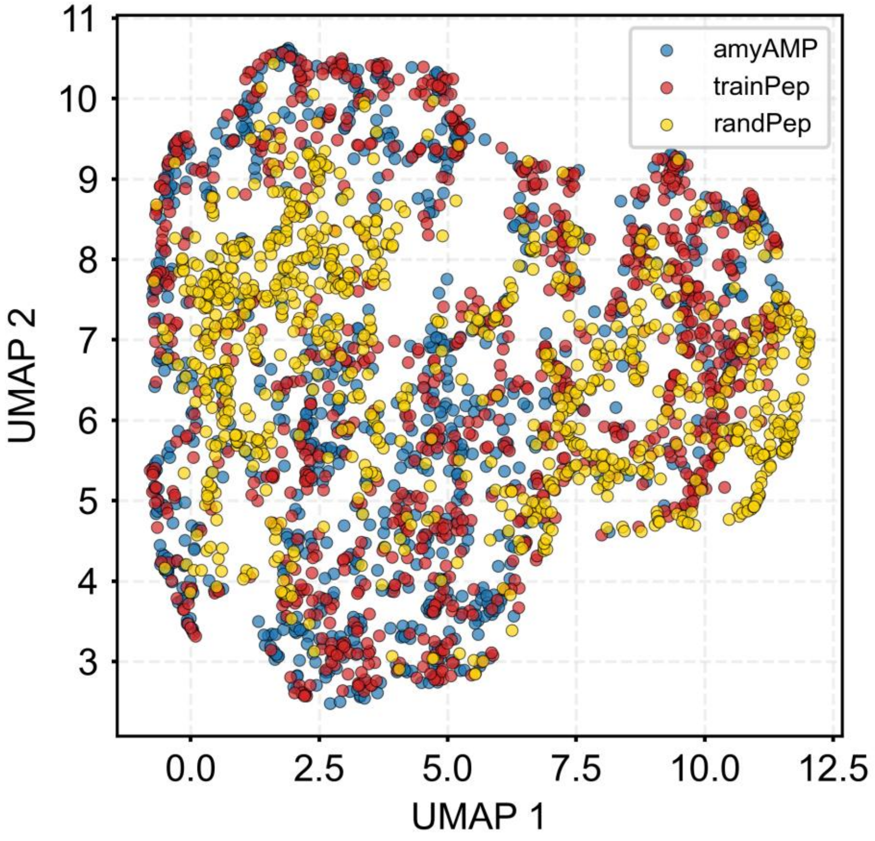


**Figure S2. UMAP projection of peptide sequence space based on PC6 physicochemical encoding.** Two-dimensional embedding generated using Uniform Manifold Approximation and Projection (UMAP) applied to peptide representations derived from the PC6 encoding scheme. The projection compares amyAMP-generated peptides (amyAMP), the combined training dataset containing antimicrobial and amyloid peptides (trainPep), and a reference set of randomly generated peptides (RandPep). The amyAMP-generated peptides largely overlap with the trainPep distribution, whereas RandPep sequences occupy a distinct and distant region. This distribution indicates that the amyAMP model successfully captures the physicochemical characteristics of the training peptide space while generating sequences distinct from random peptide compositions.

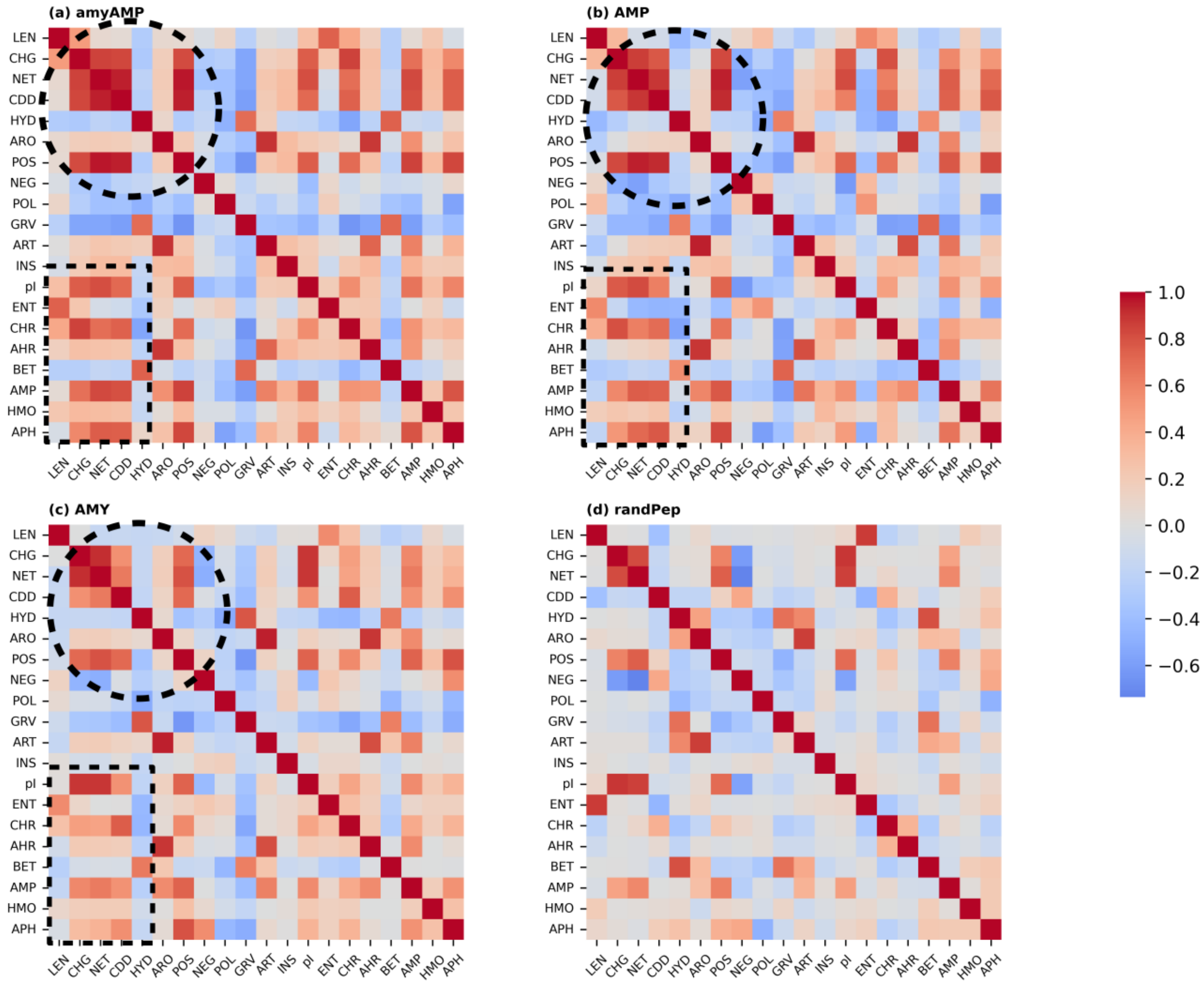


**Figure S3. Correlation analysis of physicochemical properties across peptide datasets.** Pairwise correlations among key physicochemical properties, including sequence length, charge, net charge, charge density, hydrophobic fraction, aromatic fraction, positive fraction, negative fraction, polar fraction, GRAVY, and aromaticity, were computed for peptides in four datasets: amyAMP-generated peptides, antimicrobial peptides (AMPs), amyloidogenic peptides (AMYs), and randomly generated peptides (RandPep). Full name of the abbreviations mentioned in figure are given in Table S2. The correlation patterns observed for amyAMP peptides closely resemble those present in the AMP and AMY datasets, indicating that the generated sequences preserve the intrinsic physicochemical relationships characteristic of biologically functional peptides. In contrast, these structured correlations are largely absent in the RandPep dataset, reflecting the lack of biologically relevant sequence constraints in randomly generated peptides.

**Table S2: Key physicochemical properties**

| S. No | Property Name | Abbreviation |
|---|---|---|
| 1 | length | LEN |
| 2 | charge | CHG |
| 3 | net_charge | NET |
| 4 | charge_density | CDD |
| 5 | hydrophobic_fraction | HYD |
| 6 | aromatic_fraction | ARO |
| 7 | positive_fraction | POS |
| 8 | negative_fraction | NEG |
| 9 | polar_fraction | POL |
| 10 | Gravy (grand average of hydropathy) | GRV |
| 11 | aromaticity | ART |
| 12 | instability_index | INS |
| 13 | isoelectric_point | pI |
| 14 | entropy | ENT |
| 15 | charge_hydrophobic_ratio | CHR |
| 16 | aromatic_hydrophobic_ratio | AHR |
| 17 | beta_propensity | BET |
| 18 | amp_favorable_fraction | AMP |
| 19 | hydrophobic_moment | HMO |
| 20 | amphipathicity | APH |

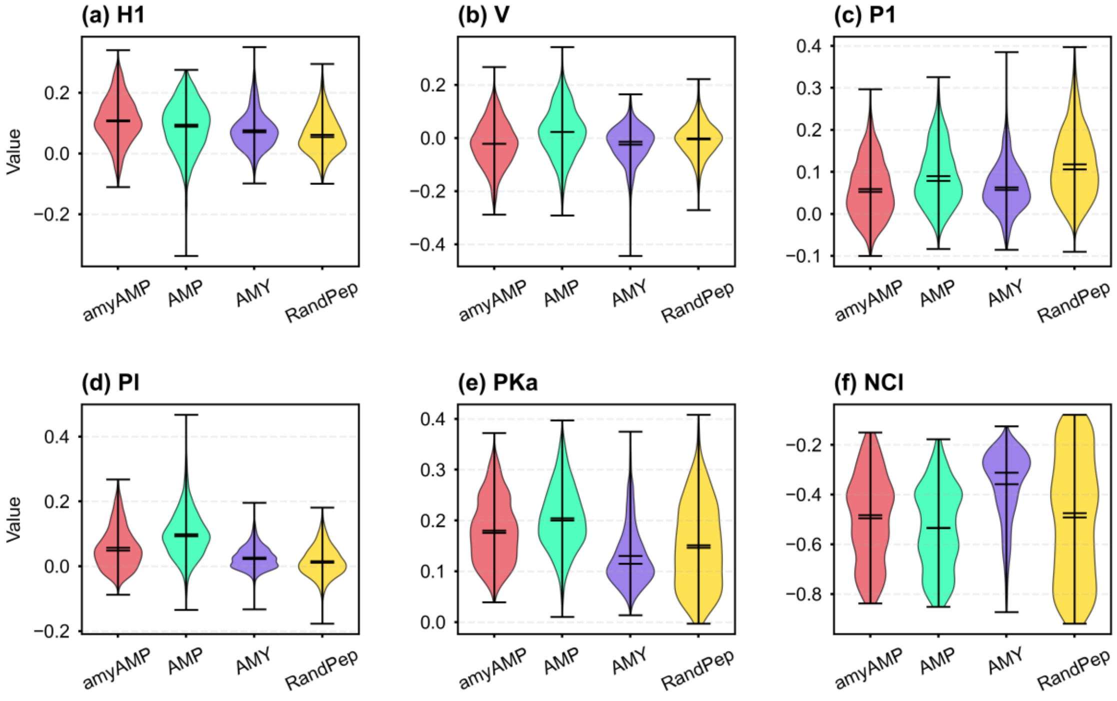


**Figure S4. Comparison of PC6 physicochemical property distributions across peptide datasets.** Violin plots illustrating the distribution of the six PC6 physicochemical properties hydrophobicity, hydrophobic moment, polarity, side-chain volume, isoelectric point, and net charge, for all peptide groups analyzed. The amyAMP-generated peptides exhibit property distributions that closely resemble those of the antimicrobial (AMPs) and amyloid (AMYs) training datasets, while clearly diverging from the random peptide control set. This comparison demonstrates that the generative model successfully captures the characteristic physicochemical landscape of functional amyloidogenic antimicrobial peptides.

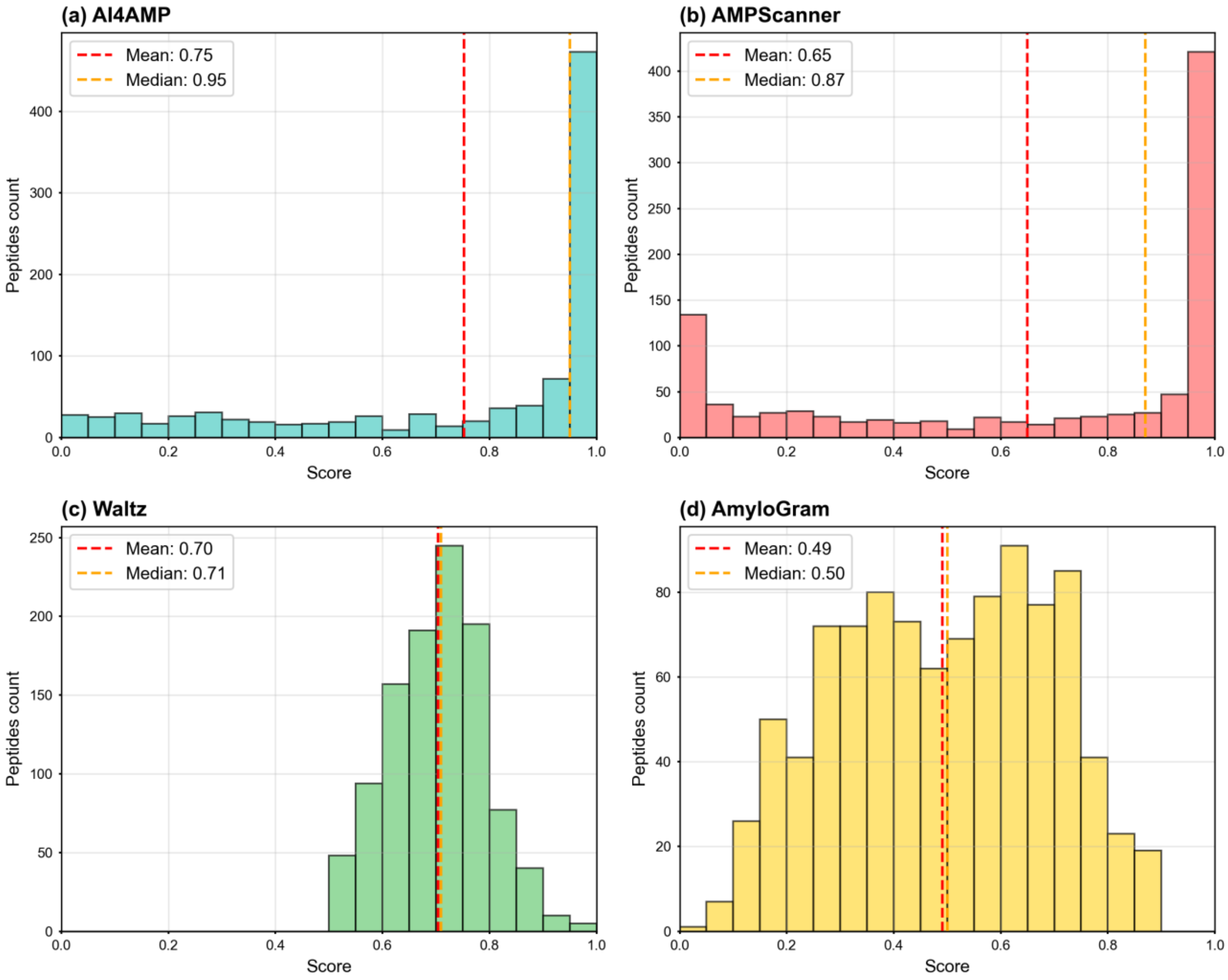


**Figure S5. Distribution of AMP scores predicted by two deep-learning models (AI4AMP and AMPScanner).** (a–b) AI4AMP predictions show that amyAMP generated peptides exhibit strongly antimicrobial character, with a mean score of 7.5 and a median of 9.5. Notably, 55.2% of the sequences fall within the very high AMP score range (0.9–1.0). (d–e) AMPScanner results similarly confirm the strong antimicrobial potential of the generated peptides, yielding a mean score of 0.65 and a median of 0.87. The pie chart further illustrates that 48.3% of the sequences are classified as very high and 9.9% as high antimicrobial probability.

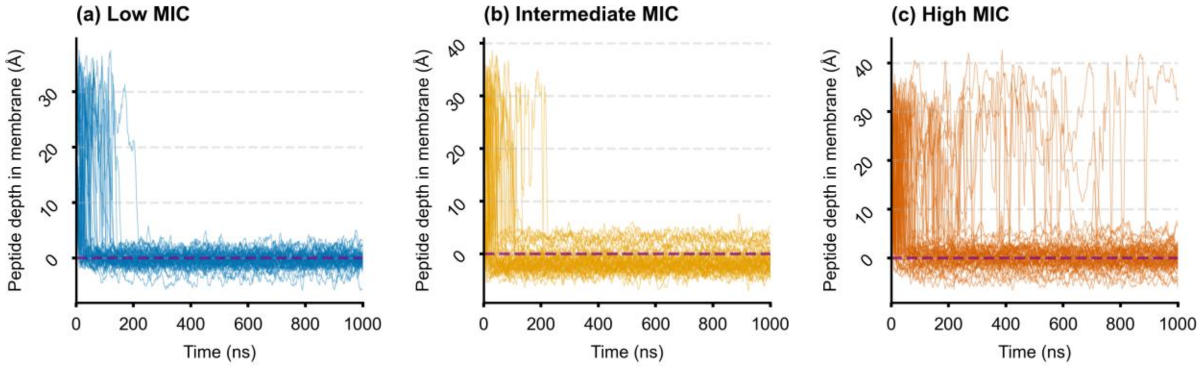


**Figure S6. Peptide insertion depth profiles for reference antimicrobial peptides grouped by activity class.** Insertion depth relative to the membrane surface is shown for peptides categorized by minimum inhibitory concentration (MIC): (a) low MIC, (b) intermediate MIC, and (c) high MIC classes. The profiles illustrate the distribution of peptide positions within the lipid bilayer during the simulations, enabling comparison of membrane insertion behavior across peptides with different antimicrobial potencies.

**Table S3: Peptide simulated in Membrane disruption simulations**

| | |
|---|---|
| High-MIC analog | SSVLISR |
| Intermediate –MIC analog | GFLSVVKKVASLIQKV |
| Low-MIC analog | RSFAKKVTKQLFTMLASYTTKNV |

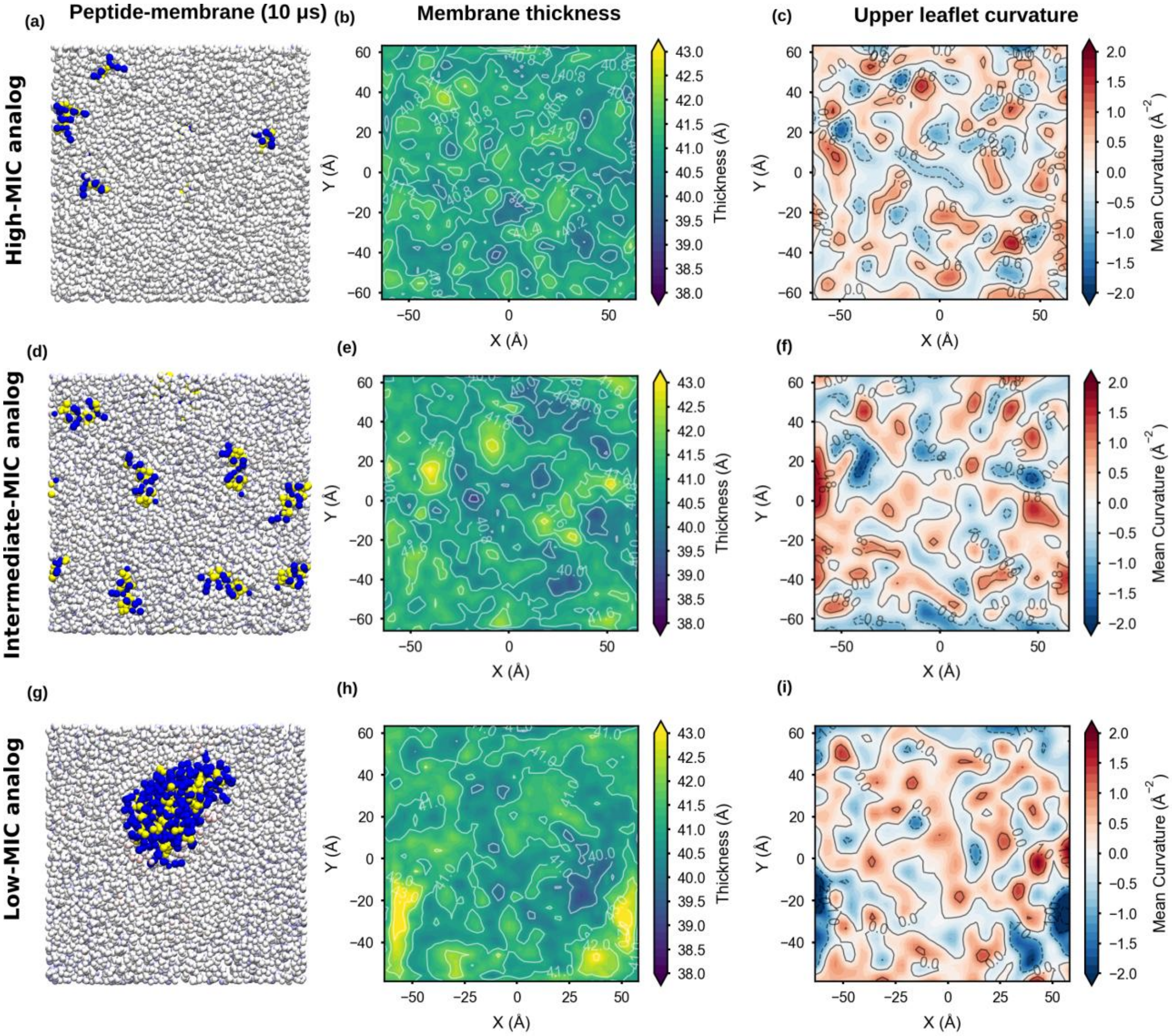


**Figure S7. Membrane disruption analysis of amyAMP peptides on a Gram-negative membrane model.** Coarse-grained molecular dynamics simulations (10 μs) of three representative amyAMP peptides, high-, intermediate-, and low-MIC analogs, interacting with a Gram-negative membrane mimic. (a, d, g) simulation snapshots at 10 μs illustrating peptide organization on the membrane surface; (b, e, h) membrane thickness maps showing local bilayer thinning; and (c, f, i) mean membrane curvature profiles reflecting peptide-induced membrane deformation. Compared with the Gram-positive membrane system (Figure 7), peptide adsorption and clustering are reduced on the Gram-negative membrane, resulting in weaker membrane thinning and smaller curvature changes, consistent with diminished peptide-membrane interactions.
